# Cell-cell communication in yeast using auxin biosynthesis and auxin responsive CRISPR transcription factors

**DOI:** 10.1101/020487

**Authors:** Arjun Khakhar, Nicholas J Bolten, Jennifer Nemhauser, Eric Klavins

**Affiliations:** University of Washington Department of Bioengineering; University of Washington Department of Electrical Engineering; University of Washington Department of Biology

## Abstract

A true engineering framework for synthetic multicellular systems requires a programmable means of cell-cell communication. Such a communication system would enable complex behaviors, such as pattern formation, division of labor in synthetic microbial communities, and improved modularity in synthetic circuits. However, it remains challenging to build synthetic cellular communication systems in eukaryotes due to a lack of molecular modules that are orthogonal to the host machinery, easy to reconfigure, and scalable. Here, we present a novel cell-to-cell communication system in *Saccharomyces cerevisiae* (yeast) based on CRISPR transcription factors and the plant hormone auxin that exhibits several of these features. Specifically, we engineered a *sender* strain of yeast that converts indole-3-acetamide (IAM) into auxin via the enzyme iaaH from *Agrobacterium tumefaciens*. To sense auxin and regulate transcription in a *receiver* strain, we engineered a reconfigurable library of auxin degradable CRISPR transcription factors (ADCTFs). Auxin-induced degradation is achieved through fusion of an auxin sensitive degron (from IAA co-repressors) to the CRISPR TF and co-expression with an auxin F-box protein. Mirroring the tunability of auxin perception in plants, our family of ADCTFs exhibits a broad range of auxin sensitivities. We characterized the kinetics and steady state behavior of the sender and receiver independently, and in co-cultures where both cell types were exposed to IAM. In the presence of IAM, auxin is produced by the sender cell and triggers de-activation of reporter expression in the receiver cell. The result is an orthogonal, rewireable, tunable, and arguably scalable cell-cell communication system for yeast and other eukaryotic cells.

## Introduction

Multicellular systems in nature are capable of incredible feats of distributed computation and self-organization. Examples range from division of labor in filamentous algae^1^, to the exquisite sensitivity of the adaptive immune system^2^, to morphogenesis and development of tissues, organs, and multicellular organisms. Computer scientists have shown that cells are in principle capable of computing a wide variety of functions^3^, generating complex morphologies^4^, and of making decisions^5,6^. Experimentally, synthetic multicellular systems have been built to regulate populations^7^, synchronize oscillations^8^, form patterns^9–11^, implement logic functions through distributed computation^3^, and cooperate to solve problems^12^. However, a scalable suite of cell-cell communication modules has yet to emerge. In particular, in *Saccharomyces cerevisiae*, strategies that use components of native signal transduction pathways can lead to crosstalk and undesirable phenotypes such as growth arrest^9,13,14^. Such systems are not obviously portable to other eukaryotes, are difficult to reprogram, and require significant changes to the host cell to function correctly^7^. Here, we describe progress toward building an engineering framework for yeast cell-cell communication that is orthogonal to yeast (and many other eukaryotic cells except plants^15^), modular, and tunable.

Orthogonality is crucial for rationally engineering cell-cell communication. Auxin, a plant hormone, does not have measurable effects on laboratory strains of yeast^16,17^ when grown in standard conditions. Our receiver cells use elements of the *Arabidopsis thaliana* auxin signaling pathway. Auxin regulates plant development via a system of transcriptional corepressors, the Aux/IAA proteins (referred to as IAAs), which are degraded in the presence of the molecule auxin. Auxin stabilizes the interaction between the degron domain of an IAA and an auxin F-box protein (AFB). The result is the degradation of the IAA via polyubiquitination^18^. The IAAs exhibit a range of degradation rates and sensitivities to auxin that are determined, in part, by the sequence of their degron domains and in part by the AFB^16,19^. The degradation dynamics of a large range of auxin degrons with multiple AFBs have been previously studied and thoroughly characterized in yeast^16^. By using this signaling modality as the basis for our communication system, we avoid using any native yeast (or mammalian) signal transduction machinery associated with adverse phenotypes^7^. Additionally, the primary components of the pathway, AFBs and IAAs, have been shown to function in several different mammalian cells^15^, suggesting that our system may be broadly portable.

To maximize modularity, we engineered auxin responsiveness into CRISPR transcription factors (CTFs). CTFs consist of a nuclease null Cas9 protein (dCas9) fused to a transcriptional effector domain. The dCas9 can be programmed to target a locus by coexpressing a small guide RNA (gRNA) that has complementarity to the target locus at a site that is adjacent to an ‘NGG’ sequence, called the PAM sequence. This strategy, as demonstrated by Farzadfard et al.^20^ and Qi et al.^21^, has the benefit of modularity through easily programmable specificity: dCas9 requires only the expression of a new guide RNA for retargeting. In contrast, zinc finger or TAL DNA binding domains require the design of a new protein for each target^22,23^. These characteristics make CTFs an ideal candidate for signal reception and processing, as they can be targeted to any promoter in the genome that has a suitable PAM site^20^, can either activate or repress gene expression, and can be layered to form more complex networks^23,24^. In the present case, CTFs fused to the VP64 strong activator domain were targeted to a promoter upstream of GFP. In addition, these CTFs were fused to Aux/IAA degron domains and co-expressed with AFBs thereby producing auxin-degradable CRISPR transcription factors, or ADCTFs. An ADCTF is thus a modular, coupled sensor-actuator, which should allow cell-to-cell communication to be easily rewired to arbitrary outputs.

Signal production and reception in cell-cell communication is ideally tunable to achieve a broad range of sensitivities and other functions. To implement and tune auxin production in the sender, we integrated the bacterial iaaH gene from *Agrobacterium tumefaciens* into yeast under the control of a constitutive promoter (GPD). Upon the addition indole-3-acetamide (IAM), sender cells produced a strong enough auxin signal to affect gene regulation via the ADCTFs in co-cultured receiver cells. The concentration of auxin produced can be tuned via the concentration of exogenously added IAM. For increased tunability, we developed a library of ADCTFs, each with a different degron and/or degron location, that displays a range of degradation kinetics and sensitivities to auxin. The sensitivity of the ADCTFs can be further tuned by the selection of either the AFB2 or TIR1 F-box as the auxin receptor. Thus, components of the ADCTFs, the auxin degron, and the transcriptional effector domain can all be swapped to obtain, respectively, a range of auxin sensitivities, and repression versus activation.

In summary, the combination of sender and receiver modules described here forms the foundation of an orthogonal, modular, and tunable cell-cell communication framework for yeast. We demonstrate each of these aspects of the system below by describing how the senders and receivers behave in isolation, and that they can be combined in co-culture to form a simple communication channel.

## Results

### Synthetic, scalable, auxin-modulated transcription factors

To link an auxin sensor to diverse transcriptional responses and targets, we designed auxin degradable CRISPR transcription factors (ADCTFs) with three modular domains (Figure 1A). The core component of the ADCTFs is the CRISPR-based transcription factor described by Farzadfard et al^18^, wherein a deactivated Cas9 protein functions as a programmable DNA binding module. The dCas9 was fused to a transcriptional effector domain, in this case the transcriptional activator VP64, and to an IAA degron. In the presence of an AFB, ADCTFs should be ubiquitinated and degraded when exposed to auxin. We tested the ADCTFs by targeting them to activate the expression of EGFP from a minimal CYC1 promoter and observed deactivation of fluorescence upon the addition of auxin. In the absence of auxin, functional ADCTFs acted identically to controls lacking a gRNA (Figure 1B). When a functional (coexpressed with gRNA) activator ADCTF was degraded in the presence of auxin, fluorescence dropped to levels at or below the control (no gRNA) levels. Auxin dependent regulation was independent of the promoter being regulated by the ADCTF (Supplementary Figure 1B). The observed effect was also reversible: when auxin was removed from the system, reporter expression returned to its activated state (Supplementary Figure 1A).

**Figure 1.**
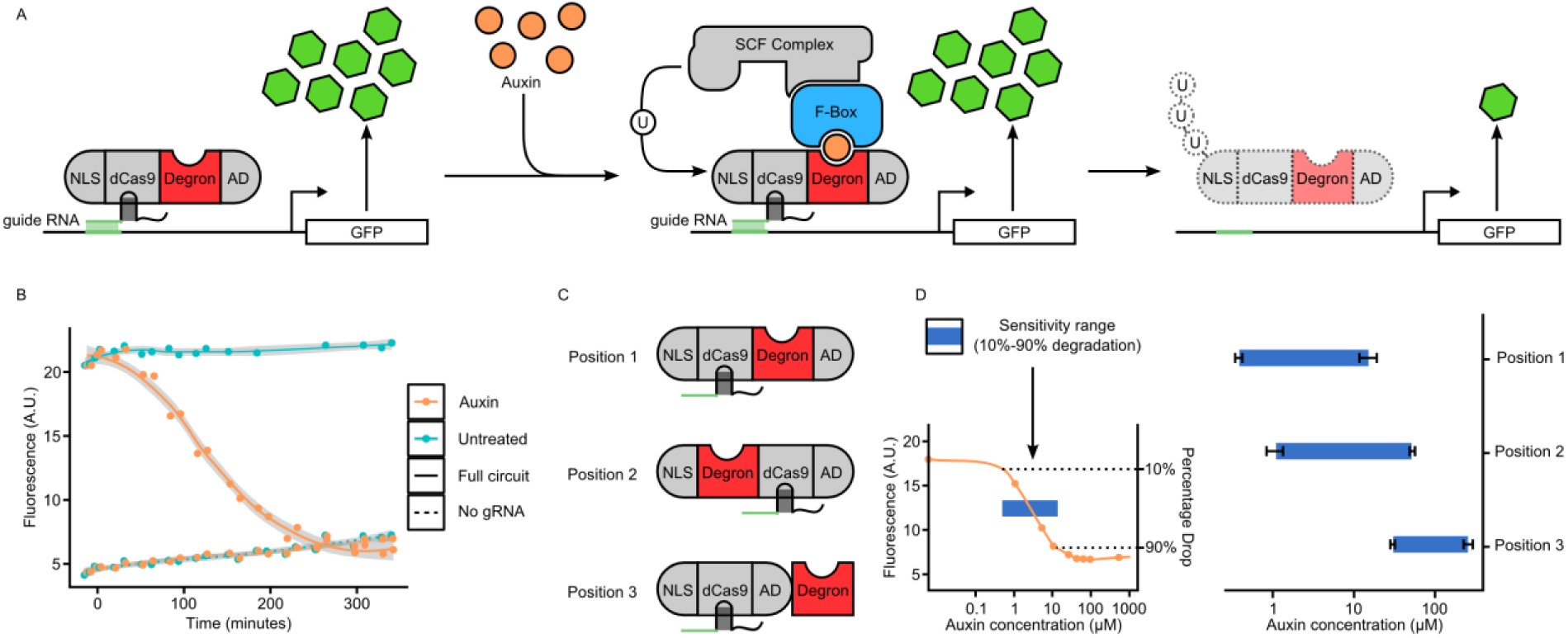
A) The ADCTF design and the molecular mechanism behind its function. An ADCTF is made up of a dCas9 protein fused to an NLS, an activation domain and an auxin sensitive degron. In the presence of auxin, the degron recruits an Auxin Sensing F-box (AFB) protein to form an SCF complex (an E3 ubiquitin ligase). The subsequent ubiquination and degradation of the ADCTF deregulates the gene targeted by the ADCTF. B) Time-lapse cytometry of ADCTF cells with a GFP-producing gRNA target following the addition of auxin or no treatment as well as with and without a guide RNA. The gray ribbon indicates the 95% confidence interval. Following treatment with auxin, the GFP level of the strain expressing gRNA dropped to basal levels (equivalent to a strain with no gRNA). C) Schematic representation of the three fusion proteins tested for the effect of degron position on ADCTF properties. D) Sensitive range characterization of the three degron position variants at steady state. Horizontal bars indicate the range of auxin concentrations between which mean steady-state fluorescence (measured via cytometry) drops from 90% of maximum to 10%. A larger sensitive range correlated with higher maximum fold changes upon induction (Supplementary Figure 2).

One design consideration for building the ADCTFs was the position of the degron within the fusion position. We hypothesized that degron position could alter accessibility to the AFB or otherwise interfere with protein folding thus modulating auxin sensitivity. We explored several possible positions for the degron relative to the other domains (Figure 1C). In all cases, the degron was flanked by flexible linkers composed of five repeats of the amino acid sequence “GS” to limit fusion-associated misfolding. Changing the position of the degron dramatically altered the sensitivity range, defined as the range of auxin concentrations between which steady-state fluorescence drops from 90% of maximum to 10% (Figure 1D). Position one is sensitive to the lowest levels of auxin, but also saturates earlier than positions two and three. Placing the degron on either side of dCas9 (positions one and two) resulted in higher auxin sensitivity than position three where the degron was placed at the C-terminal end of the fusion. The percentage drop from maximal activation upon auxin induction was directly correlated to auxin sensitivity, with position one dropping to basal levels at steady state, and positions two and three having progressively smaller effects post induction (Supplementary Fig 2). Altering the position of the degron coarsely tuned the upper and lower bounds of the sensitivity range of the ADCTF. However, since the position one variant was the most sensitive to auxin and had the highest fold change, we chose to fuse degrons in all further ADCTF variants at position one.

### Engineered ADCTF variants exhibit a broad range of auxin sensitivities and degradation kinetics

The Aux/IAA family of 29 transcriptional corepressors have been shown to exhibit a large range of degradation rates and sensitivities to auxin in yeast^16^. This range of responses to the same auxin signal is thought to result in part from the sequence of the different IAA degron domains, and in part from the varying activities different auxin-signaling F-box proteins, each showing different affinities for specific IAAs. We built a library of ADCTFs using degrons from IAA14, IAA15, and IAA17 and coexpressed them with either of two AFBs (AFB2 or TIR1). These degrons have been previously characterized as encompassing a range of auxin-induced degradation rates. In general, AFB2 promotes faster degradation of IAAs than TIR1. In addition, we included a recently characterized mutant of TIR1, TIR1-DM^25^, which has been shown to greatly accelerate auxin-induced TIR1 degradation.

All pairwise combinations of ADCTFs and F-box proteins were tested for their temporal response and dose response to auxin. Temporal responses, all performed with 30 μM auxin induction, exhibited a range of degradation kinetics that depended on both the choice of ADCTF degron and the F-box protein (Figure 2B). The kinetics, characterized by the time to 50% degradation, can be coarsely tuned by the choice of F-Box protein used, with TIR1-DM being the fastest overall, followed by AFB2 and TIR1. Within this coarse tuning, the choice of degron allowed for smaller changes in kinetics. The ADCTF with the degron from IAA15 (ADC15) seemed to have the overall fastest kinetics. The only exception to this trend was the interaction between AFB2 and ADC17, which had the fastest degradation rate. All the ADCTFs had approximately the same percentage change from maximal activation upon auxin induction at steady state. Thus, tuning kinetics by swapping F-box proteins or degrons had a minimal effect on the steady state response to auxin. Most variants dropped to approximately seventy 75% of maximal activation at steady state with a few between ten percent higher or lower than the mean (Supplementary Fig 3D). The ADCTFs exhibited varied sensitivity to auxin that depended on the combination of the degron on the ADCTF and the F-box protein. Swapping F-box proteins allowed for more coarse grain tuning of sensitivity range with TIR1 conferring the broadest sensitivity range overall and TIR1-DM conferring the narrowest (Figure 2C). Swapping degrons allows smaller changes, as was observed within the AFB2 variants wherein there is a progressively narrower sensitivity range from ADC14 to ADC17. The dynamics and steady state behavior of the ADCTFs in response to auxin correspond to the behavior of previously characterized IAA proteins, from which the degrons were taken, in yeast^14^. The only exception being the degron 17 variant, which had much slower degradation kinetics in the ADCTF context in a TIR1-DM background. This result suggests that the auxin responsive behavior may be predictably tuned by swapping degron and F-box protein variants.

**Figure 2.**
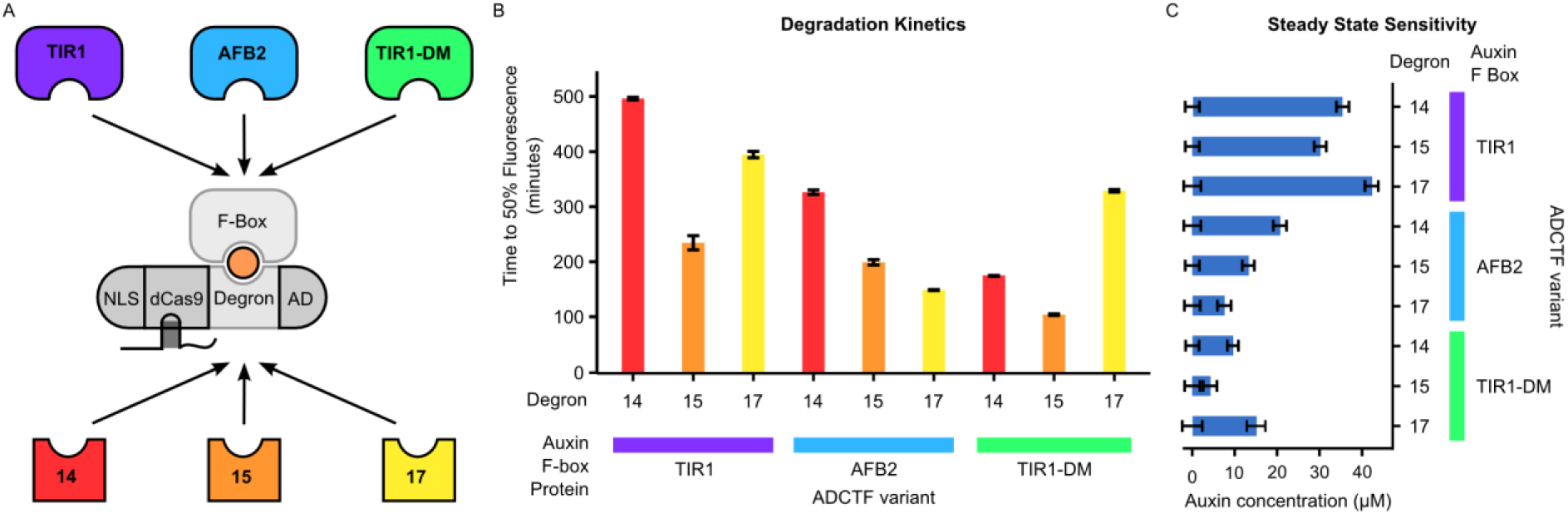
A) The ADCTF (auxin receiver) strain library was generated from all pairwise combinations of three Auxin Sensing F-box protein variants (AFB2, TIR1, TIR1-DM) and three auxin degron variants (from IAA14, IAA15 and IAA17). B) Receiver strain library degradation kinetics measured via time-lapse cytometry. The kinetics of ADCTF responses to auxin were characterized by the time at which fluorescence dropped to fifty percent of maximum: a smaller time implies a faster response. The ADCTF library displays a wide range of degradation kinetics that were modulated by both the choice of F-box protein and the auxin degron. C) Auxin sensitivity ranges for the ADCTF library. Blue bars represent the auxin sensitivity range at steady state as defined in Figure 1.

### Yeast produce tunable levels of auxin via expression of iaaH from *Agrobacterium tumefaciens*

To generate an auxin producing strain, we integrated half of the IAM pathway from *Agrobacterium tumefaciens* into yeast^26^. The IAM pathway is a two-step enzymatic process that converts tryptophan to IAM and then into auxin. The first step is via tryptophan-2-monooxygenase (iaaM, not examined here). The second step is catalyzed by indole-3-acetamide hydrolase (iaaH). To test whether yeast could produce auxin from IAM using only the second enzyme (iaaH), we integrated the iaaH gene from the *Agrobacterium tumefaciens* Ti plasmid^27^ into an auxin reporting yeast strain (Figure 3A) containing a IAA-YFP fusion protein. After adding IAM, reporter degradation rates were measured via time-lapse cytometry (Figure 3B). Upon the addition of IAM, sender strains produced an auxin response comparable to that of native auxin (Figure 3C). In addition, for a given concentration of IAM, the steady state fluorescence values converge to those of auxin (Figure 3D). There was no significant delay between the addition of IAM and the production of auxin, so the transport and production of auxin from IAM can be assumed to be faster than the reporter’s dynamics. We then investigated intercellular auxin production by coculturing the sender strain with an auxin sensor strain that could be distinguished via its mCherry signal (Figure 4A). Rather than a dose response of IAM, increasing fractions of sender cells were cocultured with sensor strains in constant amount of the auxin precursor (300 μM) to test the dependence of auxin production on sender cell concentration (Figure 4B). Greater concentrations of sender cells produced a greater auxin response in sensor cells, though both the kinetic and steady state behavior suggest that there is less auxin in the media than in sender cells.(Figure 4C, Figure 4D).

**Figure 3.**
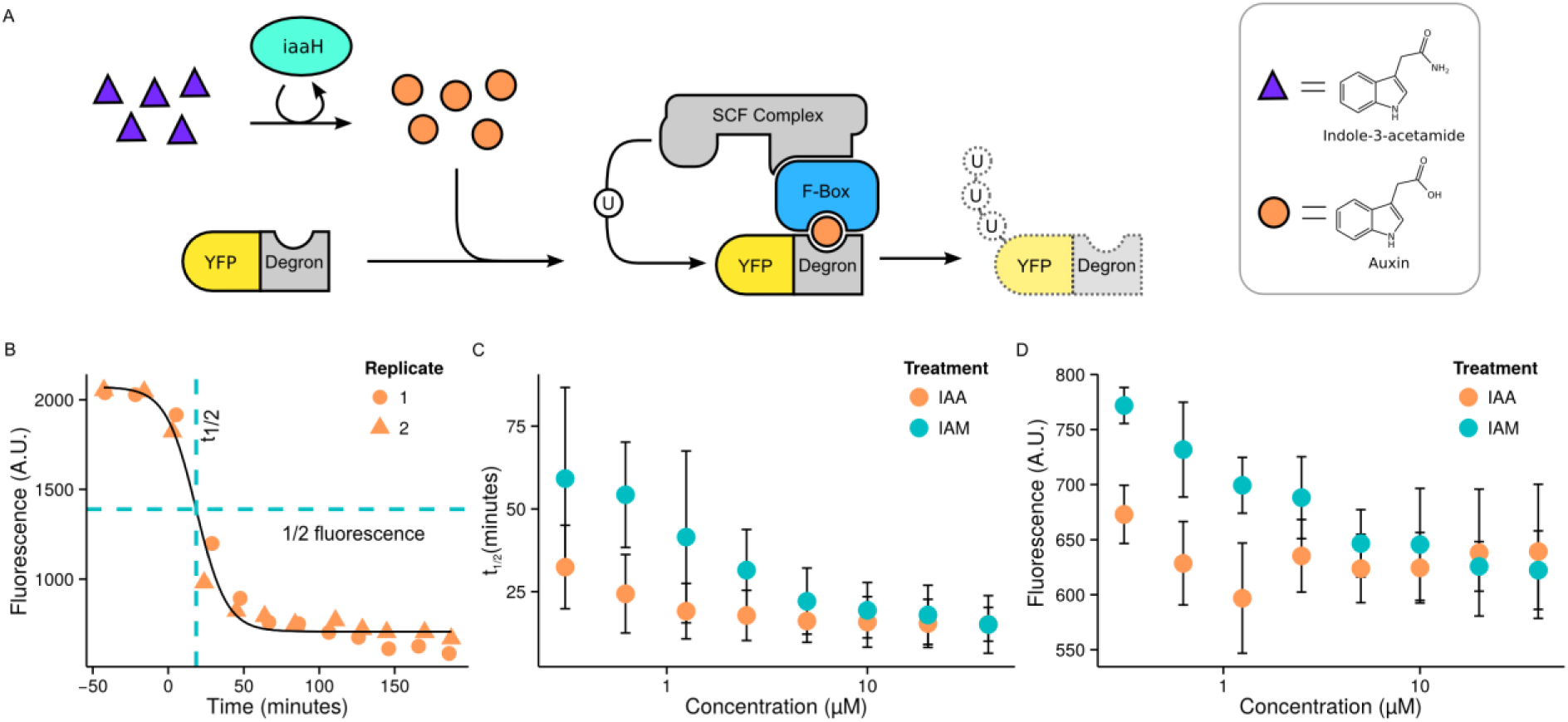
A) Auxin sender strain design. The iaaH enzyme of *Agrobacterium tumefaciens* catalyzes the conversion of indole-3-acetamide (IAM) into auxin, inducing the degradation of proteins fused to an auxin degron. The iaaH enzyme (sender cells) was integrated into an auxin reporting strain (the EYFP-IAA17|AFB2 strain from *Havens et al*^16^) to test for internal auxin production. B) Kinetic auxin response to IAM addition in sender strains. Following the addition of IAM, the fluorescence of sender cells decreased to basal levels. The time to half-maximal (t_1/2_) fluorescence was used to measure the rate of reporter degradation. C) Auxin-induced degradation rate in response to varying doses of either IAM or auxin. Sender cells were treated with either auxin or IAM and read at regular intervals producing time courses as in part B. Nonlinear fitting was used to generate t_1/2_ values. For a given molarity, treatment with IAM produces an auxin-induced degradation similar to, but weaker than, direct treatment with auxin. D) Steady state fluorescence in response to varying doses of either IAM or auxin taken from the same dataset as part C. As the concentration of IAM was increased, a lower steady state fluorescence was produced.

**Figure 4.**
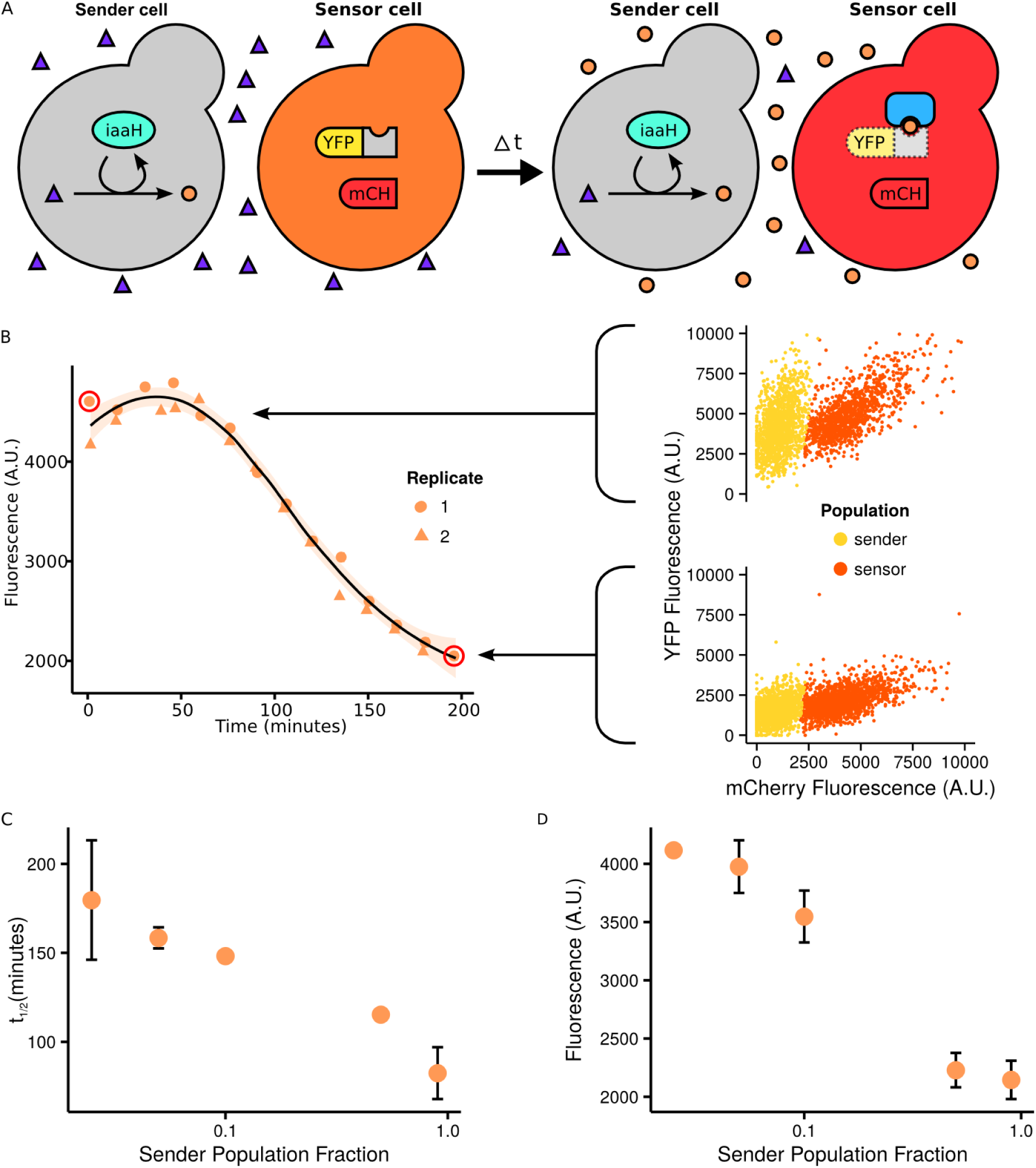
A) Sender-sensor multicellular auxin signaling strains. Sender cells are identical to those in Figure 3 and therefore produce auxin upon the addition of exogenous IAM and sense auxin production via an EYFP-IAA17 reporter. Sensor cells express an EYFP-IAA17 and TIR1 and are distinguished experimentally through the expression of mCherry. In coculture, IAM diffuses into sender cells where it is converted into diffusible auxin that then degrades EYFP-degron proteins in either the sender or sensor cell types. B) Auxin-induced degradation of EYFP-IAA17 in sensor cells cocultured with sender cells in 300 μM IAM. Data for sensor cells can be separated from sender cells via their mCherry signal. The line represents a LOESS fit and the light orange ribbon represents a 95% confidence interval of the fit. C) Sender cell fraction dose response. Each fraction had the same volume, so a larger fraction indicates a larger concentration of sender cells in coculture. As the sender cell population increases, the degradation rates decreases. D) Steady state fluorescence in response to varying doses of either IAM or auxin taken from the same dataset as part C. As the concentration of sender cells was increased, a lower steady state fluorescence was produced, flatting out at around a 50:50 split.

### Sender cells produce a tunable auxin response in receiver cells

Sender and receiver cells were cocultured in different ratios to measure the effect of sender cell concentration on auxin signal production. Senders constitutively express iaaH and the receivers expressed an activating ADCTF and a gRNA targeting a minimal CYC1 promoter driving EGFP (Figure 5A). After adding a saturating amount of the IAM and growing the coculture overnight, we observed a reduction in gene activation in the receiver strain comparable to direct addition of auxin (Figure 5B). Three different receiver strains with a range of responses to auxin were tested with the sender strain. All the receiver strains produced an auxin response and behaviors were consistent to those observed via the direct addition of auxin, suggesting that the sender module is compatible with any ADCTF-based receiver module (Supplementary Figure 4). In addition, a 10% fraction of sender cells is sufficient to a significant change in fluorescence in receiver cell at steady state and a 50% fraction produces a nearly saturating signal (Figure 5C, Figure 5D).

**Figure 5.**
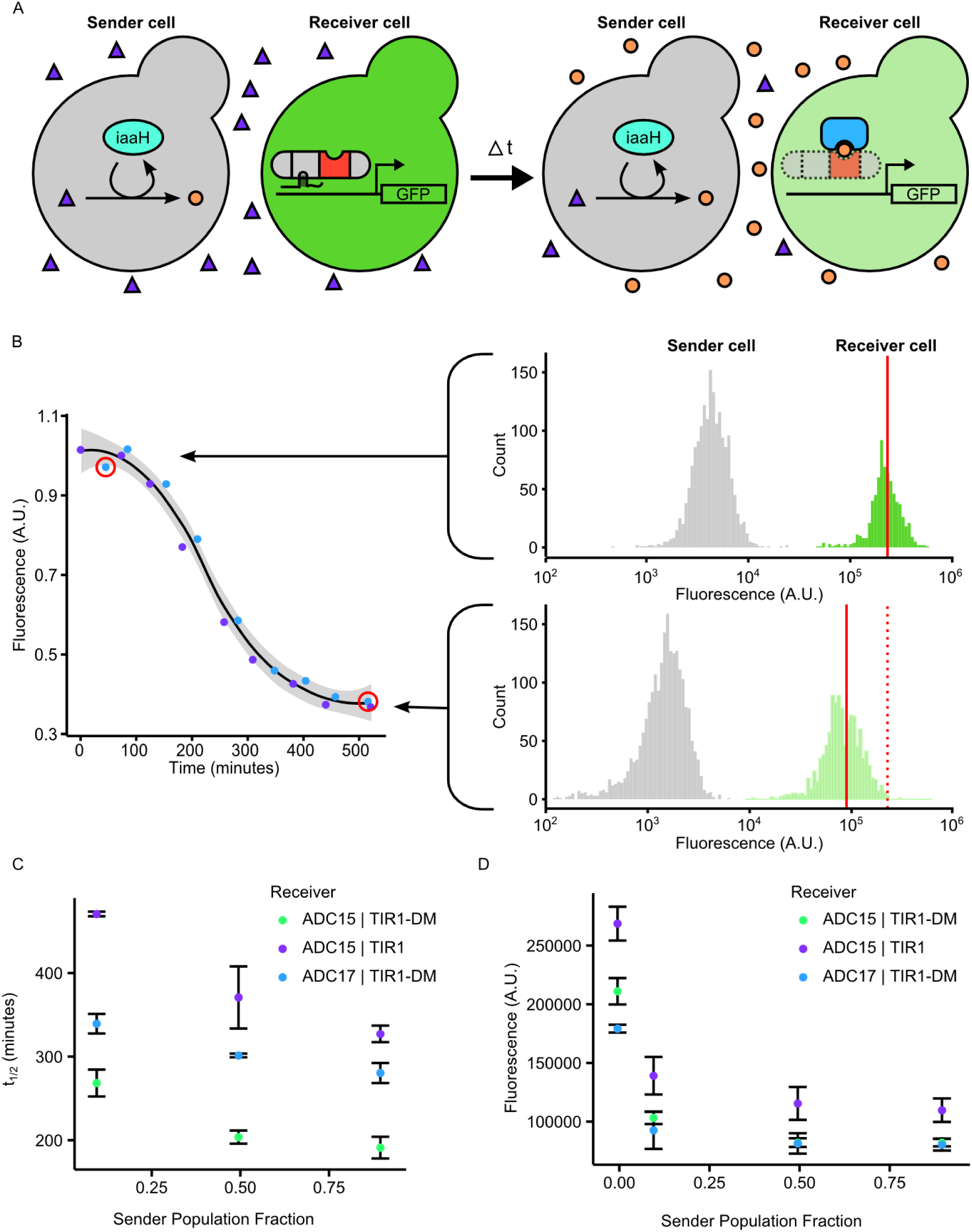
A) Coculture of sender and receiver strains. Sender cells convert IAM into auxin that then diffuses out of sender cells and into receiver cells where it causes the degradation of ADCTFs, producing a drop in fluorescence. B) Time course data for two replicates (shown in blue and purple dots) of a coculture of equal concentrations of sender and receiver cells is plotted on the left. The line represents a LOESS fit and the gray ribbon represents a 95% confidence interval of the fit. On the right, histograms display distinct populations of sender (gray, left) and receiver (green, right) cells. In the presence of sender cells treated with IAM, receiver cells dropped in fluorescence over time. As in figures 3 and 4, sender cells express also an EYFP-IAA and AFB auxin reporter and therefore also show a decrease in fluorescence. Without IAM, receiver cells did not show a significant decrease in fluorescence. C) Degradation rates (measured as t_1/2_) in receiver strains in response to sender cell concentration. As the fraction of sender cells increased, there is a more dramatic auxin effect in receiver cells that saturates at approximately even fractions of send to receive.

## Discussion

Our system is based on a signal transduction modality that is unique to plants and so is orthogonal to native yeast signal transduction pathways, as well as to mammalian cells^15^. The simplicity of the system will hopefully allow it to be ported to other contexts, such as mammalian cells. The ADCTF library allows the generation of a range of responses to the same auxin signal, and can in principle be connected to any gene of interest, or to another synthetic gene circuit. Additionally, auxin production levels can also be tuned by titrating in different amounts of IAM. It may also be possible to tune the diffusivity of auxin in yeast^26^, or to harness the sequestration and turnover pathways of auxin found in plants. Our approach of detecting small molecules via F-Box mediated degradation of a transcription factor is potentially scalable as there are other plant hormones such as jasmonate that use a very similar signaling pathway^28^. Current work involves building on these characteristics to produce more complex multicellular behaviors. For example, feedback systems can be built through regulation of the iaaH gene via the ADCTFs. More generally, our results form the basis platform for implementing distributed decision making, pattern formation, and other complex cell-to-cell communication based multicellular behaviors.

## Methods

### Strain construction

Building off the work of Farzadfard et al^20^, the reporter is a yeast-enhanced green fluorescent protein driven by a truncated CYC1 promoter. This reporter was integrated at the URA3 locus in the genome of the W303-1A ADE2 strain of *Saccharomyces cerevisiae* and this reporter strain was used as the parent for all ADCTF strains. All gRNA was driven by an ADH1 promoter driven construct that consists of a gRNA flanked on each side by a hammerhead and an HDV ribozyme, facilitating expression from an RNA polymerase II promoter. All the gRNA constructs were integrated at the HIS3 locus. AFB2, TIR1 and TIR1-DM were integrated, respectively, at the LEU2 loci, and were driven by the GPD promoter. The ADCTFs were constructed by fusing an SV40 nuclear localization tag, a VP64 activation domain, and an auxin degron to a nuclease null version of the Cas9 protein from *Streptococcus pyogenes*. The auxin degron used for all characterization, unless otherwise mentioned, was the t1 truncation of the degron from IAA17 from *Arabidopsis* that was characterized previously to have the fastest speed of degradation in the presence of AFB2 degradation machinery^16^. The other degrons used were the domain two regions from IAA14 and IAA15. The ADCTF is driven by a beta-estradiol inducible version of the GAL1 promoter integrated at the TRP locus in the genome in all strains^29^. The iaaH gene was amplified via PCR from the Ti plasmid of *Agrobacterium tumefaciens* and cloned via the Gateway^TM^ method into a single-integrating HIS3 plasmid behind the strong TDH3 promoter. The integrating plasmid cassette was produced via digestion of the plasmid by PmeI and integrated into an auxin reporter strain via a standard lithium acetate transformation method^30^.

### Cytometry

All cytometry measurements were acquired with an Accuri C6 cytometer with attached CSampler apparatus using 488 nm and 640 nm excitation lasers and a 533 nm (FL-1: YFP/GFP) emission filter. Experiments involving time course data were taken during log phase via the following preparation: 16 hours of overnight growth in synthetic complete medium in a 30°C shaker incubator followed by 1:100 dilution into fresh, room-temperature medium. After 5 hours of growth under the same incubation conditions, 100 μL aliquots were read periodically until the completion of the experiment. For experiments involving steady state behavior, cultures were grown overnight, then diluted down in the morning 1:100 in fresh media and grown for 5 hours to log phase. They were then induced and allowed to grow for between five and twenty four hours depending on the experiment and then read on the cytometer. Cytometry data were analyzed using custom R scripts and the flowCore^31^ package using the following steps: (1) gating for the yeast population, (2) gating for separate sending / receiving strains via the yellow (GFP) and red (mCherry) channels, and the generation of mean fluorescence values.

